# Breaking Apart Contact Networks with Vaccination

**DOI:** 10.1101/2020.03.01.971630

**Authors:** Gianrocco Lazzari, Marcel Salathé

## Abstract

Infectious diseases can cause large disease outbreaks due to their transmission potential from one individual to the next. Vaccination is an effective way of cutting off possible chains of transmission, thereby mitigating the outbreak potential of a disease in a population. From a contact network perspective, vaccination effectively removes nodes from the network, thereby breaking apart the contact network into a much smaller network of susceptible individuals on which the disease can spread. Here, we look at the continuum of small world networks to random networks, and find that vaccination breaks apart networks in ways that can dramatically influence the maximum outbreak size. In particular, after the removal of a constant number of nodes (representing vaccination coverage), the more clustered small world networks more readily fall apart into many disjoint and small susceptible sub-networks, thus preventing large outbreaks, while more random networks remain largely connected even after node removal through vaccination. We further develop a model of social mixing that moves small world networks closer to the random regime, thereby facilitating larger disease outbreaks after vaccination. Our results show that even when vaccination is entirely random, social mixing can lead to contact network structures that strongly influence outbreak sizes. We find the largest effects to be in the regime of relatively high vaccination coverages of around 80%, where despite vaccination being random, outbreak sizes can vary by a factor of 20.

## Introduction

The spread of infectious diseases remains a central public health issue in the 21st century. On the one hand, emerging or re-emerging diseases with no known vaccines pose a fundamental threat, and pandemics of such diseases remain on the list of potentially catastrophic events for humanity^1, 2^. On the other hand, even vaccine-preventable diseases continue to cause substantial morbidity and mortality, for two main reasons: vaccines are generally not perfectly protective^3^–5, and a vaccination coverage of 100% is rarely achieved^6^. Immunological issues such as limited vaccine efficacy, vaccine effectiveness, extent and duration of vaccine immunogenicity contribute to an imperfect protection, and remain under active investigation for improvements. Societal issues such as limited access to vaccines, as well as medical and personal reasons that prevent individuals from getting vaccinated contribute to an incomplete coverage^7^.

Despite these issues, vaccination has substantially reduced the burden of many diseases in general, and childhood diseases in particular^8^, 9. However, some vaccine preventable diseases have been making worrying comebacks in recent years. The case of measles is particularly concerning, for numerous reasons. First, measles is one of the most infectious agents known to humans, with a basic reproductive number *R*_0_ anywhere between 12 and 18^10^. Second, measles does not only cause substantial morbidity and mortality, but has recently also been shown to diminish previously acquired immune memory of other pathogens^11^, 12. Third, the measles vaccine is one of the most efficacious and affordable vaccines, providing life-long immunity in 97% of people who have received two doses^13^, 14. Because of these factors, the WHO and other health organizations recommend^15^ a vaccination coverage of 90-95% for two routine doses of measles-containing vaccines, and most WHO member states have committed to achieving these goals. However, by 2015, the global two-dose vaccine coverage was only 61%, with high variance between countries^15^. Even in high-income countries such as those in Europe, only a few countries have achieved the coverage goal. Concerningly, the number of countries who have achieved the target has declined recently, from 14 countries in 2007 to 4 countries in 2017^16^. In early 2019, the WHO declared vaccine hesitancy to be one of the top global health issues^1^.

Interestingly, however, countries with similar vaccination coverages show markedly different patterns with respect to the number of measles cases experienced (fig. 1). For example, Canada and Switzerland have almost identical vaccine coverages, but the yearly number of measles cases per capita differ by an order of magnitude. Similarly, Germany reports an almost identical coverage than the US, but has almost an order of magnitude more measles cases per capita than the US. For what reasons could similar vaccine coverages lead to large differences in relative outbreak sizes? Some hypotheses have been put forward. For example, even with similar vaccination coverage, the risk of large outbreaks can vary if unvaccinated individuals are clustered^17^. If, for example, 10% of the population is not vaccinated, and those 10% live close to each other (geographically and socially), outbreaks will likely be larger than if those 10% are more randomly distributed in the population. In the former case, the protective effect of herd immunity is larger than in the later case of clusters of unvaccinated individuals. Such a clustering phenomenon has been argued to be a likely contributor to recent outbreaks^17^. Another hypothesis is that the speed emerging outbreaks are being tackled can vary greatly from one country to the next. In the US, measles outbreaks are treated with extreme urgency and even relatively small outbreaks receive substantial media coverage, something that is not observed in other countries.

**Figure 1.**
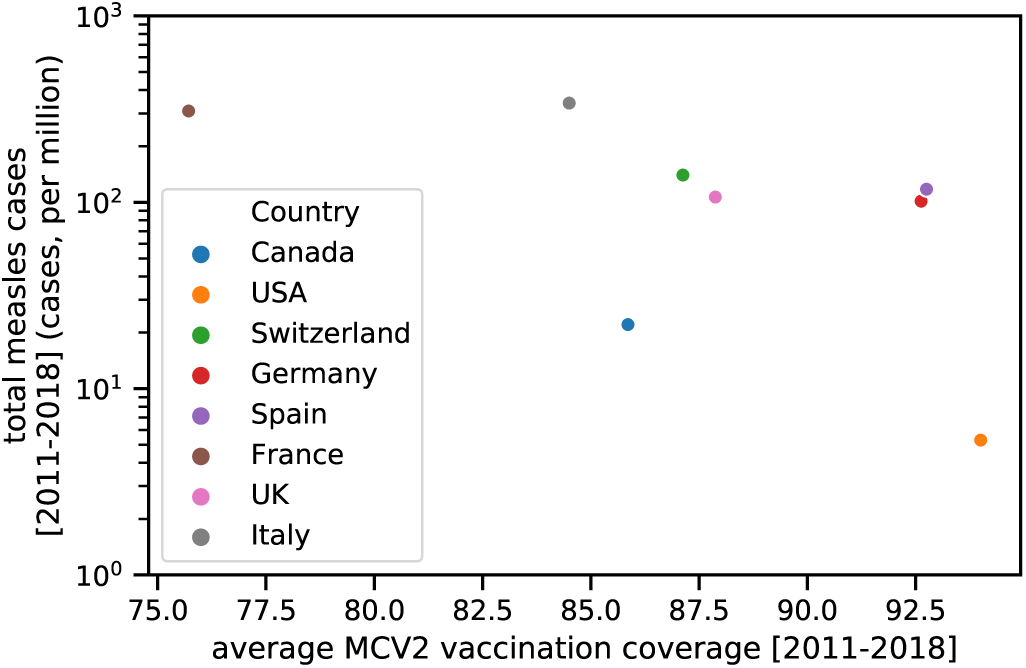
Scatter plot showing total measles cases (per million inhabitants) against average second-dose-vaccination coverage in the 2011-2018 time window, for North-American and the largest European countries. Data taken from^6, 18^

Here, we report on another phenomenon that can lead to substantially different outbreak sizes in populations with identical vaccination coverages. When large parts of a population gets vaccinated, the vast majority of possible chains of transmission is broken, thereby hampering the spread of a disease. As we will show below, the structure of the underlying contact network can greatly influence the magnitude of that effect on outbreak dynamics, and in particular on outbreak size. To do this, we will use a well-established contact network approach, where the nodes of the network represent individuals, and the edges between the nodes represent contacts along which a disease can spread. Vaccinating a node with a very effective vaccine can be thought of as removing that nodes and all its edges from the network, as no disease transmission can go through this node. When removing nodes in such a way, we are left with a much smaller and sparser network of unvaccinated nodes, on which the disease can spread. The structure of the original complete network will affect the structure of the remaining susceptible network. Indeed, with high vaccination coverages, the susceptible network will often fall apart into multiple disconnected subnetworks. This will substantial lower outbreak sizes, as the spread of the disease is confined to its network of origin, and outbreaks in a given subnetwork are limited to the size of the subnetwork. The maximal magnitude of this effect is shown to be dependent on the vaccination coverage, but given such a constant coverage, the outbreak size can differ by more than a factor of 20.

## Results

The results reported are based on network simulations, where nodes can be in one of two states, vaccinated and unvaccinated. Using measles as our infectious disease of interest, we make the simplifying assumptions that a vaccinated note is fully protected from getting infected, and that given a contact (edge) between an infected and a susceptible node, an infection is guaranteed to happen. We further ignore any timing issues with respect to incubation period and recovery times. While these assumptions do not reflect reality accurately, they are sufficiently representative to understand the worst case situation for measles, given that MCV2 status confers 97% protection, and the extremely contagious nature of the measles virus. These assumptions make stochastic disease simulations unnecessary, as the first infected node will go on to saturate the entire network of connected susceptible nodes with the disease. Thus, given multiple sub-networks (connected components) of susceptible nodes following vaccination in the complete network, and removal of all vaccinated nodes, the expected outbreak size 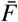 is given by 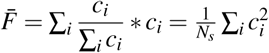, where *c*_*i*_ is the size of the i-th sub-network, and *N*_*s*_ the number of unvaccinated nodes in the networks (i.e. the sum of the size of all sub-networks). In other words, 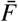 is simply the weighted sum of the sub-networks’ sizes.

In order to understand the effect of the structure of the original network on the expected outbreak size 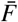, we begin with a small-world network^19^ of size *N* = 1000, and an initial rewiring probability of *p*. We then vaccinate a fraction *V* of the nodes, meaning that *V* represents the MCV2 vaccination coverage in our model. The vaccinate nodes are subsequently removed from the network, and the remaining *N*_*s*_ = (1−*V*)*N* susceptible nodes may then form multiple disconnected sub-networks, whose sizes determines the expected outbreak size 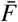 as indicated above. In order to understand the effect of the network structure on the expected outbreak size, we are calculating the outbreak size for different rewiring probabilities *p*, and different vaccination coverages *V*. Beginning from a rewiring probability *p* = 0.001, we explore increasing *p* values up to 0.8. Thus, starting from highly modular small-worlds network structures, we move increasingly towards random networks by increasing *p*, thereby lowering the modularity of the networks. Importantly, rewiring keeps the number of nodes and edges in the networks constant, making comparisons more meaningful. Figure 2 shows the effect of increasing rewiring on the size of the largest connected component of unvaccinated sub-networks (which dominates the expected outbreak size 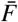 given its calculation above). Overall, less modular networks are likelier to retain a large connected component after node removal than more modular networks.

**Figure 2.**
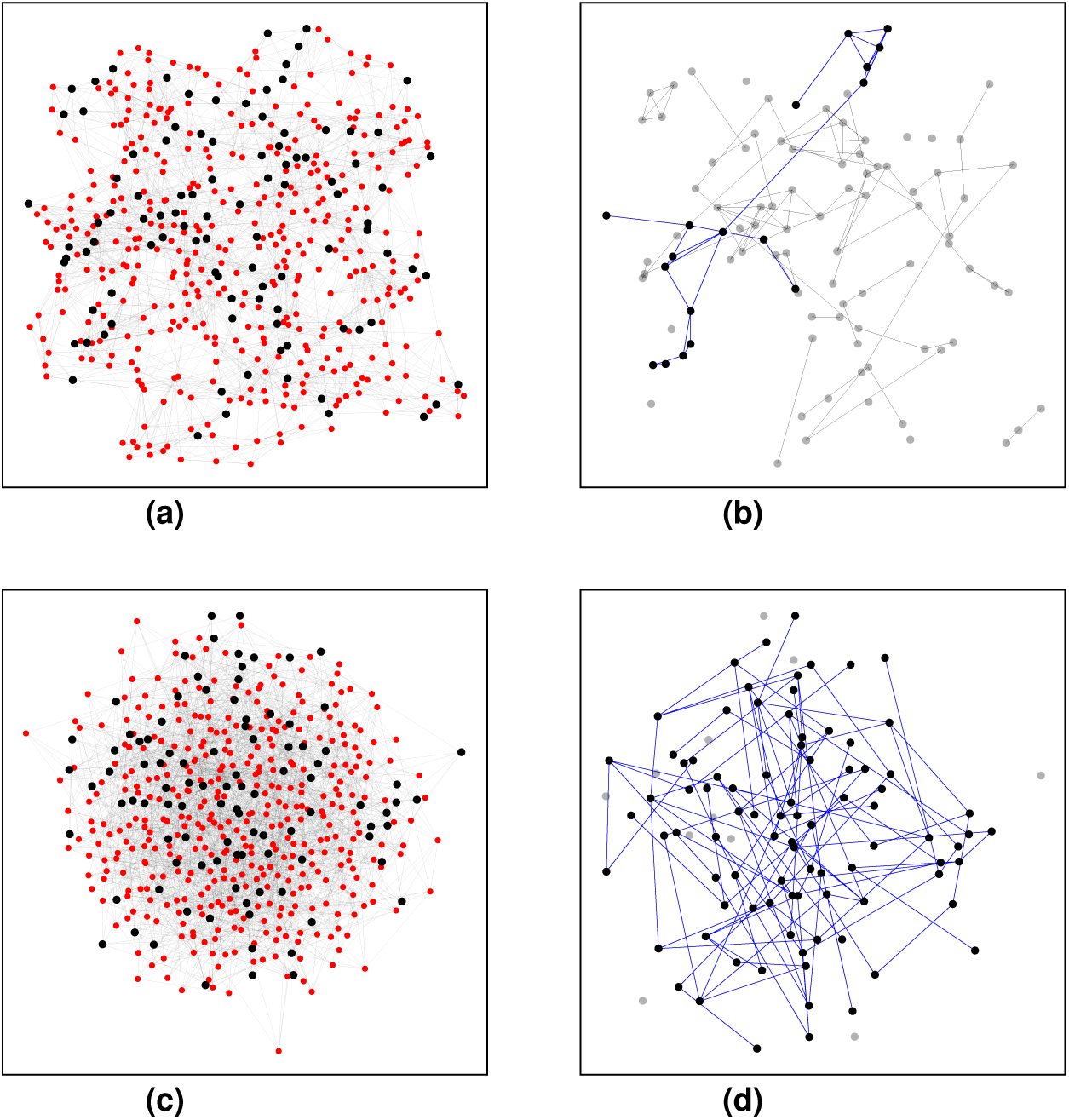
Graphs of equal size and similar structure break apart differently after random removal of 80% of all nodes. Two cases are shown, starting from the same Watts-Strogatz network (*N* = 500 and *k* = 10), but with different rewiring values: *p* = 0.1 in panel (a) with degree coefficient of variation *CV* = 0.096, and *p* = 0.8 in panel (c) with degree coefficient of variation *CV* = 0.214. In the right column, the largest connected component of the resulting graph after node removal (vaccination) is highlighted with blue edges. For top row, the expected outbreak size 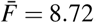, for bottom row 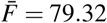.

Even though the difference in connectedness of the unvaccinated networks may appear visually subtle, as in Figure 2, its effect can nevertheless be quite consequential in terms of expected outbreak size 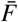. Figure 3a shows the effect of increasing rewiring on the expected outbreak size, for vaccination coverages *V* = 0.5, 0.6, 0.7, 0.8, and 0.9. While rewiring has initially little effect, we start to see noticeable effects at around *p* =0.01, initially for lower vaccination rates only, and later for higher vaccination rates as well. For each of the vaccination coverages, we can observe a transition from outbreak sizes that are far below the maximum possible outbreak sizes (as indicated by the horizontal lines in Figure 3a), approaching the maximum value with increasing rewiring. This transition spans at least an order of magnitude under all vaccination coverages, highlighting the magnitude of the effect. Overall, this demonstrates that rewiring changes the original network structure in such a way that the breaking apart of the network through vaccination-driven removal of nodes strongly influences the expected outbreak size.

**Figure 3.**
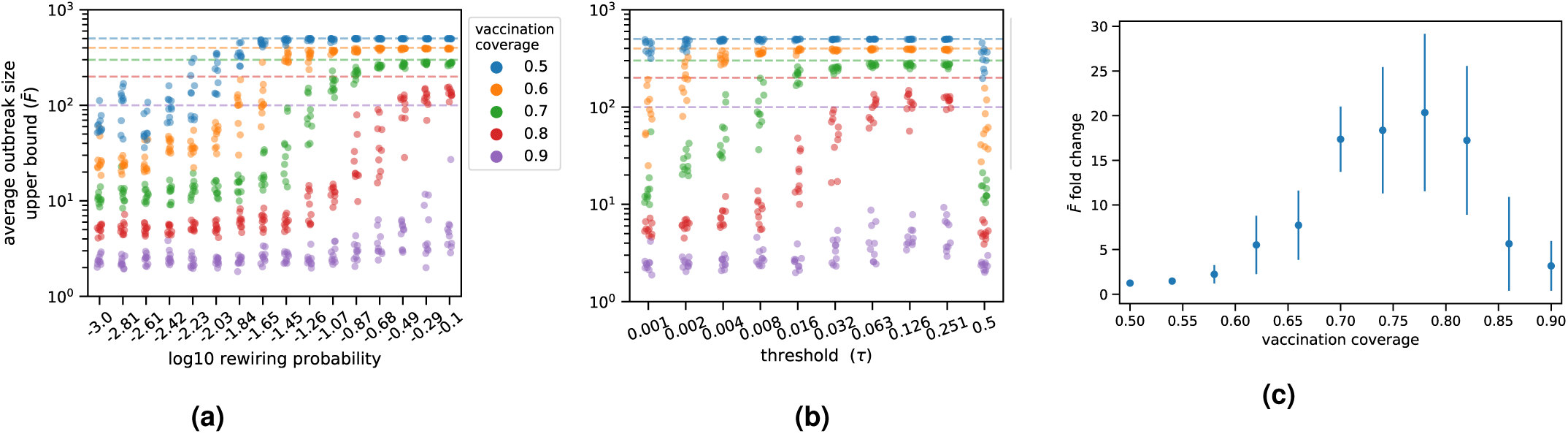
(a) Upper bound of outbreak sizes 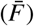, as a function of the rewiring probability *p* in the Watts-Strogatz (WS) model. 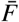 measured after running the social dynamics algorithm, as a function of the social distance threshold *τ* (see `Methods`). Each dot corresponds to a single simulations run, with 10 runs for each value of *τ* and vaccination coverage. Dashed lines in panels (a) and (b) represent the fraction of unvaccinated nodes, i.e. the theoretical maximum outbreak size. (c) Fold change of 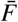, defined as the ratio between its highest value 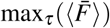 and its value at lowest threshold 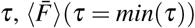, after taking mean over 10 simulation runs, for each value of *τ*. In all simulations WS graphs with *N* = 1000 and *k* = 10 were used; *p* = 0.01 in (b) and (c).

We next explore a social model that may drive the rewiring process. Social contacts may change over time for a number of reasons, and while previous infectious disease models with vaccination have focused on social dynamics due to vaccination opinions^20^, we focus here on social dynamics that are entirely independent of vaccination. To begin, we assign a random social status *s* between 0 and 1 to each node, and then rewire edges assortatively, i.e. in such a way as to implement a similarity-seeking behavior of the nodes (see Methods for detail). We measure the strength of the similarity-seeking rewiring with *τ*, which captures the threshold of dissimilarity, above which nodes seek to change their contacts to more similar nodes (with respect to social status *s*). Once the network reaches a stable equilibrium, nodes are vaccinated at random, given vaccination coverage *V*. Thus, the social dynamics in this model are independent of vaccination, and vaccination is completely random. Figure 3b shows the effect of the dissimilarity threshold *τ* on the expected outbreak size with varying vaccination coverages. We observe that the dynamics are similar to the ones described in Figure 3a. We further quantify the difference *τ* can make, given a vaccination coverage *V*, by calculating the ratio between the expected outbreak size 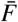 (the minimal value), and the value of *τ* where 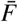 is maximal for the given vaccination coverage. Notably, at the minimal value *τ* = 0.001, there are barely any rewirings, because the desire for similarity (or rather the dislike of dissimilarity) is so great that nodes cannot find suitable similar nodes. This value thus represents largely unmodified small-worlds network. Therefore, the calculated ratio quantifies the maximum strength of the effect of social dynamics. As can be seen in Figure 3c, this ratio can reach values of up to around 20, especially at vaccination coverage around *V* ∼ 0.8. In other words, depending on the structure of the network due to social dynamics, outbreak sizes can differ by a factor of 20, even though the vaccination coverages are the same, and vaccination is at random. Importantly, these effect do not appear to be captured well by modularity - outbreak sizes can vary considerably in the range *τ* < 0.03 even though the modularity of the networks is roughly the same (see Figure 4, right panel).

**Figure 4.**
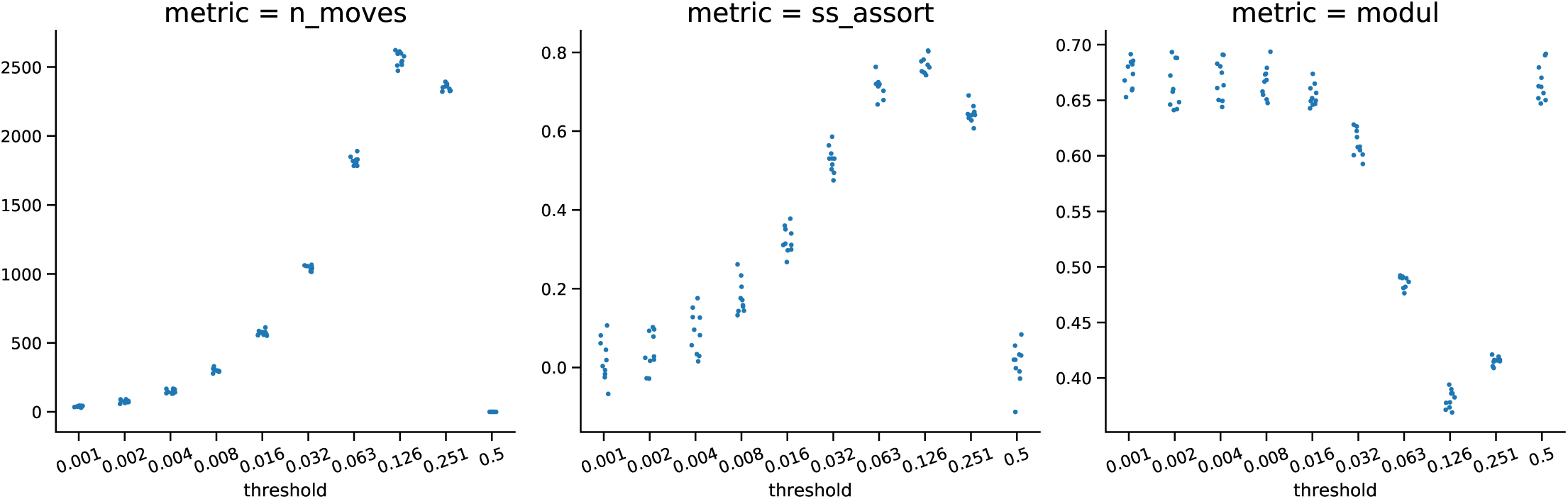
Effect of social dynamics algorithm on network properties, as a function of threshold *τ* (log scale). Number of actual edge rewirings performed (left panel), assortativity of nodes’ ‘social status’ (central panel), and network modularity (right panel). As in figure 3, dots corresponds to a single simulations run (10 simulation runs for each value of *τ*).

Finally, we explore the structural dynamics of social changes depending on the dissimilarity threshold *τ*. Low values of *τ* mean that nodes are generally seeking to connect to other, more similar nodes, but finding other nodes is challenging, given the very low dissimilarity threshold. Thus, the number of overall rewirings is low, as seen in Figure 4 (left panel). As *τ* is increasing, nodes are less likely to seek new connections, but when they do, they are more likely to find them due to the higher dissimilarity threshold. Thus, increasing *τ* leads to more rewirings. At a certain level of *τ*, the dynamics reverses, and rewirings become more rare: with increasing dissimilarity thresholds, nodes have little desire to seek out new connections. These overall dynamics of rewiring have a direct impact on the assortativity with respect to the social status *s*, and on the modularity of the network. As the rewirings are increasing, assortativity is increasing (as nodes are seeking, and finding, more similar nodes to connect to), and modularity is decreasing due to the random structural nature of the rewire (note that while the rewiring process itself is not random, but based on the value of *s*, the structural effect is nevertheless random in nature, because the values of *s* have initially been assigned randomly to nodes). This effect eventually weakens again, when *τ* becomes so high as to prevent most nodes from seeking to rewire in the first place.

## Discussion

Vaccination is a powerful tool to curb the spread of infectious diseases in human contact networks because of its ability to break apart potential transmission chains. Given sufficiently high vaccination coverage, vaccination does not only break apart transmission chains, but has the potential to break apart a large contact network into many sub-networks, therefore substantially lowering the maximum possible size of an outbreak. We showed here that the original network structure influences the sub-network structure in ways that can have very strong effects on expected outbreak sizes. In some cases, we observed a 20-fold difference of expected outbreak size, despite identical vaccination coverages.

We started from the observation that the number of measles cases per capita can differ substantially among countries even if they have very similar vaccination coverages. In particular, as can be seen in Figure 1, the number of measles cases per capita on the North American continent are roughly an order of magnitude lower than in European countries with similar vaccination coverages. While there may be multiple reasons for this, we suggest that social dynamics influencing contact network structures, as shown here, can also play a role. For example, some evidence points to higher social segregation in the US compared to Europe^21^–23. It is plausible that higher social segregation will manifest itself in higher network modularity. The lowering of network modularity is the main topological reason why vaccination would fail to break apart the network into many disjoint subnetworks. In other words, in a network with weakly inter-connected communities (and higher modularity), the same level of vaccination will more likely disconnect communities from each other than in a network with strongly inter-connected communities (and lower modularity), thereby reducing the expected outbreak size in the former.

Previous models have associated social clustering with higher probabilities of vaccine-preventable disease outbreaks^17^. Such models are generally based on the assumption of a vaccine decision-making process^20^, whereby vaccination is clustered in the network due to individuals’ beliefs about vaccination, or other personal views that are correlated with vaccine decision-making. In contrast, our model strictly assumes a random distribution of vaccination, and thus describes a different phenomenon. Given that both effects are likely to be in play in reality, it will be interesting to see how these two phenomena interact in future work.

## Methods

We generated and manipulated the networks using the `networkx` python library. In particular, we used the `community.greedy_modularity_communities`^24^ and `community.modularity` functions to compute the graphs’ modularity^25^. The Watts-Strogatz networks used for the social dynamics model (fig. 4, 3b and 3c) were generated with rewiring probability *p* = 0.01 and number of initial nearest neighbors *k* = 10. Each node was assigned a random variable (*s*, its ‘social status’), uniformly distributed in [0, 1]. Then we introduced a circular distance in the social-status space between node *n*_1_ and node *n*_2_, defined as: *d*(*s*_1_,−*s*_2_) ≡ *min*(|*s*_1_−*s*_2_|, 1−|*s*_1_ − *s*_2_|). The distance takes therefore values in the range [0, 1*/*2]. We then run our social dynamics algorithm, summarized hereby: For each edge in the graph (say (*n*_1_, *n*_2_) ∈ *E*(*G*)):

1. decide if the ‘social connection’ between the two nodes *n*_1_ and *n*_2_ is too weak, based on a global threshold *τ*: *d*(*s*_1_, *s*_2_) ≥ *τ*
2. if yes, pick at random one of the two nodes (*n*_*old*_) linked by the edge (e.g. *n*_*old*_ = *n*_1_)
3. pick at random another node of the graph (*n*_*new*_), outside of the neighborhood of *n*_*old*_ (*n*_*new*_ ∉ *N*_*G*_(*n*_*old*_))
4. if a new link is possible (i.e. *d*(*s*_*new*_, *s*_*old*_) < *τ*), rewire the old edge to the new contact (remove (*n*_*old*_, *n*_2_) and add (*nold, nnew*))

were *τ* is a free-parameter of the model. The algorithm was stopped after the actual rewiring slows dramatically; in our case we set the max number of iterations equal to 4 times the number of edges 4\**E* = 20000, much bigger than the highest number of moves actually observed (see fig. 4, left panel). Note the algorithm preserves the total number of edges, as well as the mean degree, while it does not necessary keep the graph connected.

## Author contributions statement

G.L. performed data analysis and simulations; M.S. supervised the study; G.L. and M.S. wrote the paper.

## References

1. Ten threats to global health in 2019. https://www.who.int/news-room/feature-stories/ten-threats-to-global-health-in-2019 (2019).

2. Bill gates: deadly flu epidemic is one of biggest threats to humanity - insider. https://www.insider.com/deadly-flu-epidemic-biggest-threat-bill-gates-2018-learnings-2018-12 (2018).

3. Ward, J. I. et al. Efficacy of an acellular pertussis vaccine among adolescents and adults. New Engl. J. Medicine 353, 1555–1563 (2005).

4. La Torre, G., De Waure, C., Chiaradia, G., Mannocci, A. & Ricciardi, W. Hpv vaccine efficacy in preventing persistent cervical hpv infection: a systematic review and meta-analysis. Vaccine 25, 8352–8358 (2007).

5. Osterholm, M. T., Kelley, N. S., Sommer, A. & Belongia, E. A. Efficacy and effectiveness of influenza vaccines: a systematic review and meta-analysis. The Lancet infectious diseases 12, 36–44 (2012).

6. Who | data, statistics and graphics. https://www.who.int/immunization/monitoring_surveillance/data/en/ (2019).

7. Hill, A. B., Kilgore, C., McGlynn, M. & Jones, C. H. Improving global vaccine accessibility. Curr. opinion biotechnology 42, 67–73 (2016).

8. Rappuoli, R., Pizza, M., Del Giudice, G. & De Gregorio, E. Vaccines, new opportunities for a new society. Proc. Natl. Acad. Sci. 111, 12288–12293 (2014).

9. Peter, G. Childhood immunizations. New Engl. J. Medicine 327, 1794–1800 (1992).

10. Guerra, F. M. et al. The basic reproduction number (r0) of measles: a systematic review. The Lancet Infect. Dis. 17, e420–e428 (2017).

11. Petrova, V. N. et al. Incomplete genetic reconstitution of b cell pools contributes to prolonged immunosuppression after measles. Sci. immunology 4 (2019).

12. Mina, M. J. et al. Measles virus infection diminishes preexisting antibodies that offer protection from other pathogens. Science 366, 599–606 (2019).

13. Rosenthal, S. R. & Clements, C. J. Two-dose measles vaccination schedules. Bull. World Heal. Organ. 71, 421 (1993).

14. Demicheli, V., Rivetti, A., Debalini, M. G. & Di Pietrantonj, C. Vaccines for measles, mumps and rubella in children. Evidence-Based Child Heal. A Cochrane Rev. J. 8, 2076–2238 (2013).

15. Who position paper on measles vaccine. https://www.who.int/immunization/policy/position_papers/WHO_PP_measles_vaccine_summary_2017.pdf?ua=1 (2017).

16. Ecdc: Insufficient vaccination coverage in eu/eea fuels continued measles circulation. https://www.ecdc.europa.eu/en/news-events/ecdc-insufficient-vaccination-coverage-eueea-fuels-continued-measles-circulation (2019).

17. Salathé, M. & Bonhoeffer, S. The effect of opinion clustering on disease outbreaks. J. The Royal Soc. Interface 5, 1505–1508 (2008).

18. Who | measles and rubella surveillance data. https://www.who.int/immunization/monitoring_surveillance/burden/vpd/surveillance_type/active/measles_monthlydata/en/ (2019).

19. Watts, D. J. & Strogatz, S. H. Collective dynamics of ‘small-world’networks. Nature 393, 440 (1998).

20. Mbah, M. L. N. et al. The impact of imitation on vaccination behavior in social contact networks. PLoS computational biology 8 (2012).

21. DiPrete, T. A., Gelman, A., McCormick, T., Teitler, J. & Zheng, T. Segregation in social networks based on acquaintanceship and trust. Am. journal sociology 116, 1234–83 (2011).

22. Mossong, J. et al. Social contacts and mixing patterns relevant to the spread of infectious diseases. PLoS medicine 5 (2008).

23. Hens, N. et al. Mining social mixing patterns for infectious disease models based on a two-day population survey in belgium. BMC infectious diseases 9, 5 (2009).

24. Clauset, A., Newman, M. E. & Moore, C. Finding community structure in very large networks. Phys. review E 70, 066111 (2004).

25. Newman, M. E. Modularity and community structure in networks. Proc. national academy sciences 103, 8577–8582 (2006).

